# Harmonizing terminal deoxynucleotidyl transferase dUTP nick end labeling (TUNEL) with multiplexed iterative immunofluorescence enriches spatial contextualization of cell death

**DOI:** 10.1101/2024.09.04.611218

**Authors:** Marc S Sherman, Thomas McMahon-Skates, Lindsey S. Gaston, Joseph A Majzoub, Wolfram Goessling

## Abstract

Terminal deoxynucleotidyl transferase dUTP Nick End Labeling (TUNEL) is an essential tool for the detection of cell death in tissues. Although TUNEL is not known to be compatible with multiplexed spatial proteomic methods, harmonizing TUNEL with such methods offers the opportunity to delineate cell-type specific cell death labeling and precise spatial contextualization of cell death in complex tissues. Here we evaluated variations of the TUNEL assay for their compatibility with a multiplexed immunofluorescence method, multiple iterative labeling by antibody neodeposition (MILAN), in two different tissues and injury models for cell death, acetaminophen-induced hepatocyte necrosis and dexamethasone-induced adrenocortical apoptosis. Using a commercial Click-iT-based assay as a standard, TUNEL signal could be reliably produced independent of antigen-retrieval method, with tissue-specific minor differences in signal-to-noise. In contrast, proteinase K treatment consistently reduced or even abrogated protein antigenicity, while pressure cooker treatment consistently enhanced protein antigenicity for the targets tested. Antibody-based TUNEL protocols using pressure-cooker antigen retrieval were MILAN erasure-compatible thus enabling harmonization of TUNEL with MILAN. As many as four staining cycles could be performed without loss of subsequent TUNEL signal, while first-round TUNEL did not influence protein antigenicity in subsequent rounds. We conclude this harmonized assay performs comparably to an established commercial assay, but preserves protein antigenicity, thus enabling versatile integration with multiplexed immunofluorescence using MILAN. We anticipate this harmonized protocol will enable broad and flexible integration of TUNEL into multiplexed spatial proteomic assays, thus vastly enhancing the spatial contextualization of cell death in complex tissues.

## Introduction

The terminal deoxynucleotidyl transferase dUTP Nick End Labeling (TUNEL) assay is a mainstay method for the characterization of cell death^1,2^, encompassing both apoptotic and necrotic processes^3,4^. Among cell death assays, TUNEL is particularly important for elaborating the spatial context of cell death in *in situ* in fixed tissue^1,5^. However, this advantage is attenuated by the inability to colocalize more than 2-3 additional protein targets by immunofluorescence, thus limiting the comprehensive multiplexed exploration of spatial and mechanistic relationships between cell death and tissue context. In addition, TUNEL requires the dedicated one-time use of a slide specimen, which can be limiting particularly with rare clinical specimens. Whether TUNEL is compatible with any of the modern multiplexed spatial proteomic approaches has not been investigated.

The past decade has seen multiple new or refined multiplexed protein labeling techniques that offer sequential or simultaneous staining of 20-80 protein targets in one tissue specimen. These include iterative indirect immunofluorescence imaging (4i)^6^, in cyclic immunofluorescence (CyCIF)^7,8^, multiple iterative labeling by antibody neodeposition (MILAN)^9,10^, the Co-detection by indexing (CODEX) platform^11,12^, and imaging mass spectrometry^13^. These methods share an antibody-based strategy to achieve spatial specificity of protein targets, with multiplicity enabled by photobleaching, reversible DNA barcoding, chemical erasure, or spatially resolved mass spectrometry.

Like TUNEL, the MILAN method^9,10^ is natively performed on formalin-fixed paraffin embedded (FFPE) sections as routinely processed in clinical pathology laboratories without the need for a special antibody conjugation or bleaching apparatus. Antibody erasure is accomplished by 66°C incubation of specimens in 2-mercaptopethanol with sodium dodecyl sulfate (2ME/SDS). This efficient erasure may even be antigen-retrieving, thus enabling tissue specimens to be preserved for many cycles of conventional immunofluorescence^9^. Additionally, after staining, the MILAN method is compatible with −20°C storage of slides for extended periods, thereby conserving precious tissue resources for later usage.

Because TUNEL is often performed using antibody-based detection of modified uridines, it is plausible that TUNEL staining is compatible with 2ME/SDS erasure. TUNEL erasure might then enable iterative immunofluorescence on the same sections, greatly enhancing the spatial contextualization of cell death and enabling preservation of precious clinical samples. However, TUNEL involves several potentially irreversible steps that differ from conventional immunofluorescence, including proteinase K (proK) treatment, fixation, and terminal deoxynucleotidyl transferase-mediated incorporation of a non-canonical nucleotide. It is also possible that idiosyncrasies of the MILAN protocol could influence or alter TUNEL reactivity. A different set of wash and antibody dilution buffers are used in MILAN compared to conventional immunofluorescence, and, in addition, the erasure step entails repeated 2-ME/SDS treatments which abolishes detection of some antigens^9^. The compatibility of MILAN and TUNEL is therefore unknown.

Here, we investigated differences in tissue processing in the MILAN and TUNEL protocols to delineate whether these methods could be harmonized on the same specimens. We sought to identify steps in both protocols that permanently altered the slides, establish whether there is flexibility in the sequence of these procedures, and specify any limitations or advantages to the choice of antigen retrieval method. Because TUNEL is known to stain cells that have undergone either necrosis or apoptosis with different morphological patterns^3,4^, we established reference standard murine models for both mechanisms of cell death. For necrosis, acetaminophen toxicity in the liver yields robust and recognizable hepatocyte death^14^. For apoptosis, we induced adrenocortical cell death through corticosteroid treatment^15^. These biological references enabled rational optimization toward a harmonized MILAN-TUNEL protocol which will improve our understanding of the tissue environment of cell death while enabling preservation of precious clinical specimens.

## Results

### Antibody-based TUNEL is erasable in 2-ME/SDS

The core potential benefit of harmonizing MILAN and TUNEL is erasure of TUNEL signal, allowing for restaining of the same section (Fig. 1A). The erasure step of MILAN involves tissue-preserving treatment of FFPE slide specimens with 2-mercaptoethanol and SDS (2-ME/SDS) at 66°C, which has been shown to remove most primary and secondary antibodies^9^. To test whether such an approach might enable erasure in the context of TUNEL, we first optimized an antibody-based readout of TUNEL^2,4^ that could test 2-ME/SDS-based erasure and serve as a foundation for further optimization.

**Figure 1.**
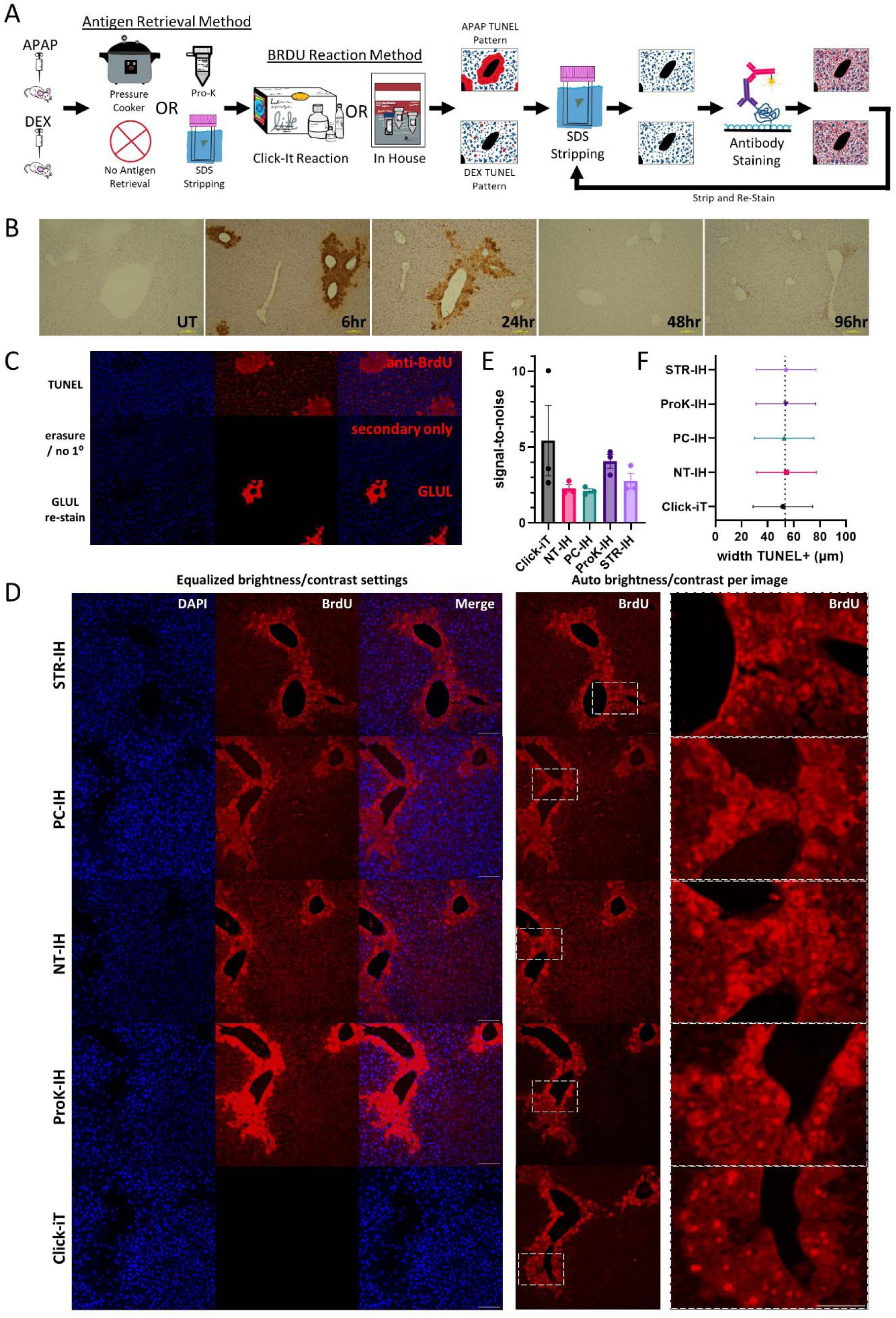
Optimizing the TUNEL protocol for compatibility with multiplexed iterative immunofluorescence. (A) Diagram of the experimental overview which benchmarks TUNEL using acetaminophen (APAP)-induced necrosis in the liver and dexamethasone (DEX)-induced apoptosis in the adrenal gland. Antigen retrieval methods were compared, a bespoke TUNEL assay was optimized in comparison with a Click-iT kit-based assay, and MILAN was conducted. (B) Time course of TUNEL positivity after acetaminophen hepatoxicity in mice shows using the Apoptag chromogenic assay. (C) Proof of concept TUNEL staining (top row) followed by erasure (secondary only, middle row) followed by staining of the affected area with a different primary and secondary (third row). (D) Direct and contrast/brightness equalized comparison of in-house TUNEL versus Click-iT (left 3 columns), and condition-optimized brightness contrast (4^th^ column) along with inset (5^th^ column), varying the antigen-retrieval method. (E) Signal-to-noise as measured by TUNEL+ areas to TUNEL-areas for each protocol, error bars are SEM, four images measured per biological replicate. (F) Measuring the thickness of TUNEL+ areas yields no difference between assays (ANOVA, p=0.99). Scalebars are 100µm.

We previously characterized a murine model of acetaminophen (APAP) hepatotoxicity and regeneration^14^. In this model, spatially restricted necrosis in the first 1-2 cell layers around central veins is maximal at 6 hours after APAP exposure in male mice and was readily visualized by a commercial, horseradish peroxidase-based TUNEL assay (**Fig. 1B**). The distinctive spatial pattern of TUNEL positivity in mouse livers 6 hours after APAP treatment (6h-APAP) served as the primary positive control for testing and optimizing TUNEL protocols. Evaluation of two “in-house” anti-BrdU-based TUNEL protocols (proK-IH and PC-IH) (see Methods) qualitatively matched both the commercial horseradish peroxidase-based TUNEL kit (**Fig. 1B**) and a commercial EdU/Click-iT-based TUNEL kit result (**Figure S1A-C**, top row) (see Methods). The same protocols yielded no staining when performed on liver specimens from saline-treated control mice (**Fig. S1A-C**, row 2), or where EdU (Click-iT) or BrdU (-IH assays) were omitted (**Fig. S1A-C** row 3). All three protocols yielded pan-nuclear signal in DNAse-treated specimens (**Fig. S1A-C** row 4).

To test antibody erasure, the PC-IH 6h-APAP TUNEL-treated specimens (**Fig. 1C**, top row) were de-coverslipped and incubated in 2-ME/SDS, re-stained with secondary-antibody only, and the same microscopic fields were re-imaged demonstrating complete erasure of both primary and secondary antibodies (**Fig. 1C**, middle row). We then erased again in 2-ME/SDS (to remove residual secondary antibodies) and performed primary and secondary staining of these liver samples with anti-Glul, as glutamine synthetase (Glul) is a highly specific marker for the first few cells surrounding the central vein (Zone 3). This restaining demonstrated retention of antigenicity of pericentral hepatocytes in precisely the region most potentially affected by the TUNEL labeling reaction in Zone 3. These results indicate that an antibody-based TUNEL assay is erasure-compatible with MILAN, and that the TdT reaction does not detectably diminish costaining in TUNEL-positive areas. These results motivated further efforts to validate and harmonize TUNEL and MILAN protocols.

### Preservation of TUNEL signal in necrosis and apoptosis across antigen retrieval methods

We next investigated whether differences between our TUNEL IH-PC protocol and a gold standard commercial assay (Invitrogen Click-iT Plus TUNEL Assay Kit) introduced differences in TUNEL staining. The protocols differ at several steps; among these, antigen retrieval was the largest potential difference in tissue treatment. Antigen retrieval in the commercial TUNEL Click-iT assay is achieved with proK treatment, a step in common with every commercial kit protocol we reviewed, and with most published protocols^2,4^. However, alternatives to proK-mediated antigen retrieval have been investigated with TUNEL, including heat-mediated antigen retrieval^16,17^, pressure cooking^18^, or other protease (e.g., trypsin^19^) treatments. The published MILAN protocol calls for no antigen retrieval^9,10^, as the 2-ME/SDS is thought to be intrinsically antigen-retrieving. We previously found MILAN is compatible with a pressure-cooker based antigen retrieval in TE buffer (pH=9)^20^, but not with citrate (pH=6) retrieval (sections fall off the slide during erasure; unpublished observation).

To understand how antigen retrieval affects TUNEL signal fidelity, we tested each antigen-retrieval method coupled with our IH TUNEL assay in serial sections of 6h-APAP murine livers (**Fig 1A**). We considered proK treatment, no antigen retrieval as called for in the first round of MILAN (NT), pre-treatment of tissue in the MILAN 2-ME/SDS “stripping” buffer (STR), and a pressure cooker treatment in TE buffer (pH=9) (PC) (**Fig. 1A**). The Click-iT TUNEL method was performed per manufacturer instructions which includes proK antigen retrieval. Among IH assays, proK-IH yielded the brightest absolute signal, however, all five assays yielded the characteristic TUNEL staining around central veins (**Fig. 1D**, middle column). Note the Click-iT signal was much less, leading to no visualization when using identical brightness/settings (**Fig. 1D**, BrdU column), but demonstrated excellent signal to noise (see auto-contrast, 4^th^ column, **Fig. 1D**). There were subtle differences between protocols; both proK treated samples, proK-IH and proK-Click-iT, showed more prominent nuclear staining in necrotic cells for example, and had overall better signal-to-noise than PC, STR, and NT retrieval methods (**Fig. 1E**). However, necrosis is expected to induce both nuclear and cytoplasmic TUNEL staining; we therefore measured the thickness of TUNEL positivity around the central veins as a quantitative comparison, and found that TUNEL positivity thickness did not differ between antigen retrieval strategies (p=0.97, nested 1-way ANOVA, **Fig. 1F**), indicating that all methods had sufficient signal-to-noise to yield the same spatial distribution of TUNEL.

We next tested PC-IH versus the commercial Click-iT assay in quantifying apoptosis using dexamethasone-treated mouse adrenal glands. Mice were treated for 4 days with dexamethasone (DEX) or vehicle (dexamethasone naïve, or DN) via drinking water. A third group of animals was treated with 4 days of dexamethasone as above and daily, intraperitoneal injections of cosyntropin, an ACTH (1-24) analog (DEX+COS). Both protocols demonstrated minimal TUNEL activity in DN adrenal glands and profound activation of apoptosis with DEX treatment (Fig. 2A), in agreement Wyllie and Kerr^15^ (Click-iT, DN v DEX p=0.020, t-test; PC, DN v DEX p=0.009, t-test). Both protocols also demonstrated suppression of apoptosis with co-administration of COS, although neither trend reached statistical significance (Click-iT, DEX v DEX+COS p=0.13, t-test; PC, DEX v DEX+COS p=0.06, t-test). The PC-IH protocol yielded nearly twice the number of TUNEL-positive cells in DEX and DEX+COS (**Fig. 2B,C**), as well as less pronounced suppression of apoptosis in the DEX+COS condition. However, the DN samples were nearly identical, suggesting minimal false-positive TUNEL staining using PC-IH. Signal-to-noise between positive TUNEL signal and background was equivalent between PC and Click-iT in adrenal tissue (**Fig. 2D**). A pattern that emerged between 6h-APAP and the adrenal TUNEL experiments was increased variability between samples in the Click-iT assay in comparison to PC-IH (**Fig. 1E**, **Fig. 2B-D**).

**Figure 2.**
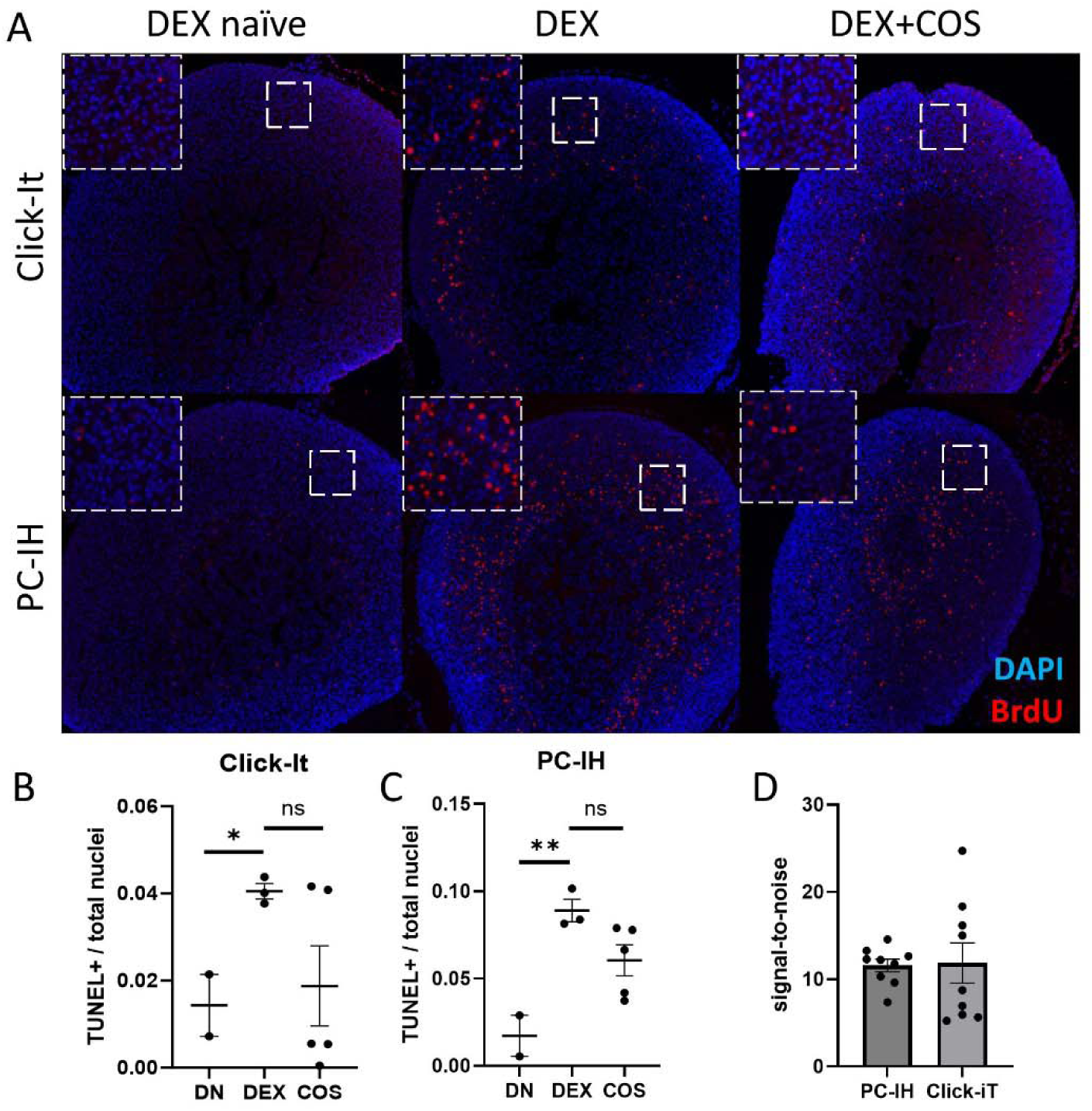
Comparison of PC-IH to Click-iT for detection of apoptosis. (A) Mice were treated with DEX, DEX+COS, or DMSO, and the adrenal glands were harvested at 4 days for Click-iT (top row) or PC-IH (bottom row) protocol TUNEL staining. Quantification of TUNEL+ nuclei over total cortical (zF+zG) nuclei by Click-iT (B) or pressure cooker (C). (D) Signal-to-noise for both protocols, where TUNEL+ signal is divided by TUNEL-DAPI spots. Scalebars are 100µm.

These data indicate that alternative antigen retrieval strategies, including pressure cooking, yield qualitatively and quantitatively similar TUNEL signal as traditional proK-based antigen retrieval in gold standard biological contexts for necrosis and apoptosis. Moreover, PC-IH exhibited reduced variability compared to a gold standard commercial TUNEL assay, highlighting its potential as a reliable and more consistent alternative antigen retrieval strategy.

### Proteinase K alters or abrogates antigenicity of multiple proteins in FFPE specimens

We next evaluated whether the antigen retrieval method affects the antigenicity of protein detection by immunofluorescence. In the same slides as Fig. 1D, we co-stained liver sections for β-catenin, a membrane bound marker of hepatocytes, and albumin, a highly expressed, secreted product of hepatocytes. Unexpectedly, we found that the Click-iT protocol completely abrogated β-catenin signal (**Fig. 3A**). Albumin staining was markedly diminished and demonstrated an aberrant, apparently nuclear-localized pattern in Click-iT slides (Fig. 3A). Normal, membrane-pattern of β-catenin was preserved in PC-IH, as was pan-hepatocyte, cytoplasmic-limited staining of albumin (**Fig. 3A**, middle row). Click-iT differs from PC by antigen-retrieval method (proK), incorporated nucleotide (EdU) and signal readout (Click-iT reaction). Therefore, proK-IH—which differs only by antigen-retrieval method—was also conducted and similarly yielded loss of β-catenin and aberrant, nuclear-pattern albumin (**Fig. 3A**, bottom row), indicating that altered antigenicity is not due to the Click-iT reaction. Both proK-IH and Click-iT share proK antigen retrieval and do not use pressure cooker retrieval. In a separate experiment, we found β-catenin was not visualized using other antigen retrieval methods (**Fig. S2**).

**Figure 3.**
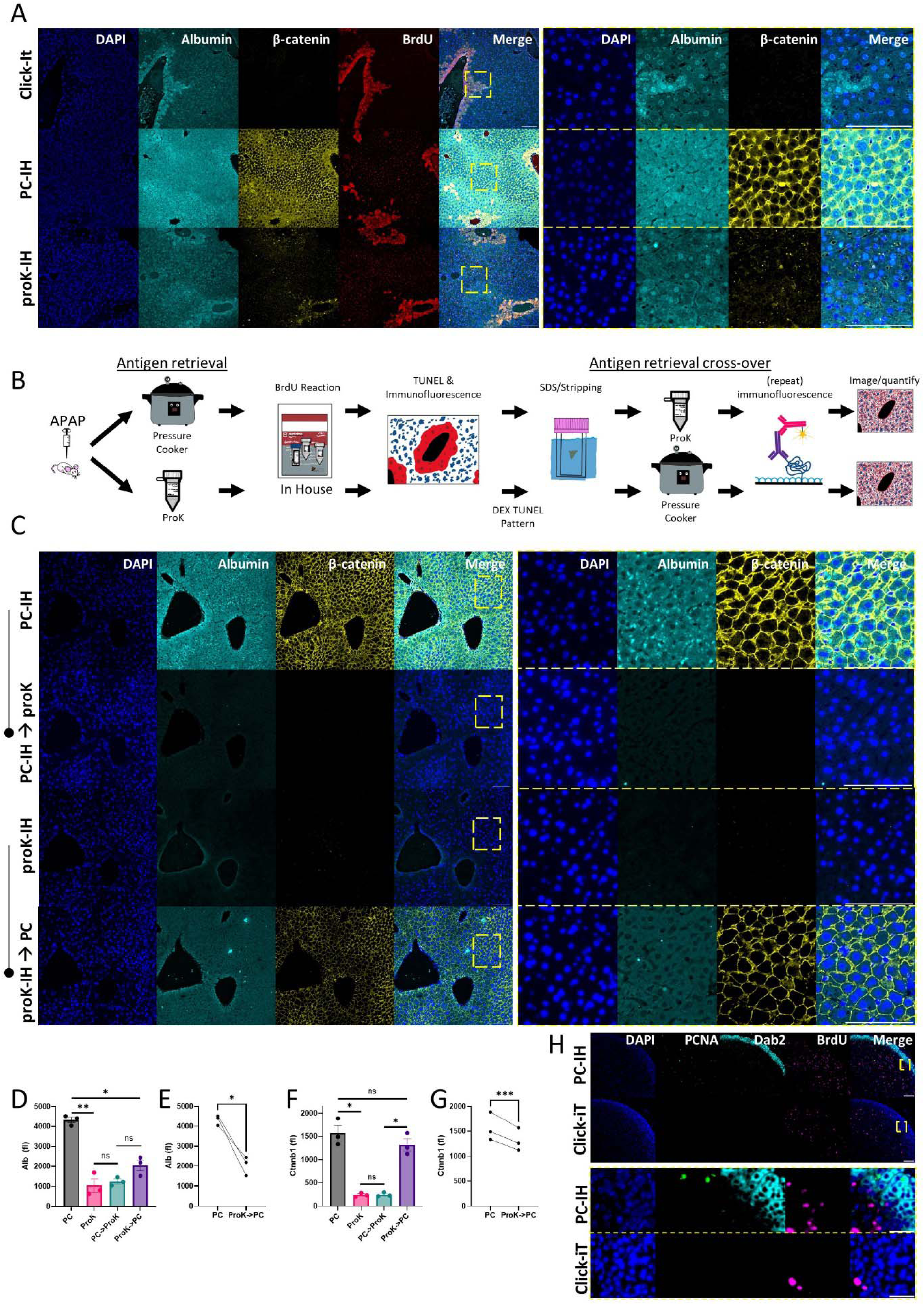
Antigen retrieval cross-over reveals proteinase K dependent signal loss, pressure cooker signal gain for multiple antigens. (A) MILAN immunofluorescence and TUNEL signal after Click-iT (top row), PC-IH (middle), or proK-IH (bottom) protocols; right columns, inset from column 5. (B) Diagramming the antigen-retrieval cross-over and restaining experiment. (C) MILAN immunofluorescence for β-catenin (yellow) and albumin (cyan) after PC-antigen retrieval (row 1), proK antigen retrieval (row 3), PC➔proK (row 2), and proK➔PC (row 4) in serial sections. (D-G) Quantification of albumin (D) and β-catenin (Ctnnb1) signal (F) across antigen retrieval methods, error bars are SEM; *=p<.05, **=p<.01, t-test. Paired analysis of PC versus proK➔PC for albumin (E) and β-catenin (F), in serial sections for the same biological replicates, *=p<.05, ***=p<.001, paired t-test, to quantify proK-dependent irreversible loss of antigen. All scalebars are 100µm.

To disentangle potential loss of signal due to digestion of antigen by proK from a gain of signal due to pressure cooker antigen-retrieval, we conducted an antigen retrieval cross-over experiment (**Fig. 3B**). Serial 6h-APAP sections in biological triplicates were antigen-retrieved and TUNEL-processed for either proK-IH or PC-IH with co-staining for β-catenin and albumin, imaged, and erased in 2-ME/SDS. PC-retrieved specimens were then treated with proK, and proK specimens were treated with PC antigen retrieval. As before, β-catenin signal was completely absent, and albumin staining was markedly diminished in proK-IH (**Fig. 3C**, rows 1 and 3).

This cross-over experiment demonstrated three independent effects of proK versus PC antigen retrieval, with minor antigen-specific differences between albumin (**Fig. 3D,E**) and β-catenin (**Fig. 3F,G**). First, PC markedly increases signal of both albumin (**Fig. 3D**) and β-catenin (Fig. 3F) compared to proK, in contrast to TUNEL signal where there is equivalence (**Fig. 2D**) or an advantage (**Fig. 1E**) of proK compared to PC depending on the tissue. Second, proK induces massive, reversible loss of signal in PC-retrieved samples (see PC versus PC➔ProK, and ProK➔PC). Third, proK induces a variable degree of irreversible antigen loss which is antigen dependent. We quantified the irreversible loss by comparing the fluorescence in serial sections treated with PC, versus proK followed by PC, yielding 53% loss for albumin (95% CI [23%-94%], Fig. 3E) or 16% loss for β-catenin (95% CI [82%-86%]) (**Fig. 3G**). We conclude that proK treatment reversibly and irreversibly diminishes both antigens to different degrees; in addition, pressure cooker treatment is required to elaborate β-catenin>albumin antigenicity.

ProK treatment—or possibly, lack of pressure cooker—also completely prevented detection of other protein immunogenicity. In the Click-iT versus PC-IH experiment in the adrenal gland, co-staining was performed for Dab2, which marks the zona glomerulosa (outer layer) of the adrenal cortex, as well as PCNA, which marks cells undergoing DNA synthesis in the cell cycle. The Click-iT commercial assay, which uses proK antigen retrieval, demonstrated complete absence of PCNA and DAB2 in adrenal sections, while both were easily detected in PC-IH (**Fig. 3H**).

These experiments highlight key advantages of PC-IH over the Click-iT commercial assay, primarily by avoiding use of proK and associated antigen loss, while retaining equivalent sensitivity for TUNEL signal and greater compatibility with conventional immunofluorescence. We therefore focused on the PC-IH protocol and harmonization with the MILAN protocol.

### Versatile integration of TUNEL into a MILAN staining series

As MILAN comprises many rounds of immunostaining, we next asked whether there were limitations to the sequence in which TUNEL (PC-IH) could be performed in the MILAN process. We sought to understand whether the TUNEL reaction itself influenced the antigenicity during concurrent or subsequent immunostaining steps, and whether MILAN or 2-ME/SDS erasure affected TUNEL staining.

First, 6h-APAP liver sections were PC antigen-retrieved, and then MILAN immunostained for albumin in the first round, without performing TUNEL (**Fig. 4A**, top row). Sections were then erased in 2-ME/SDS, and entered into the TUNEL (IH) protocol. After IH-TUNEL, we costained again for albumin (**Fig. 4A**, bottom row). TUNEL staining remained robust, despite being preceded by a round of immunostaining and 2-ME/SDS erasure. In addition, repeated staining of albumin was preserved, or slightly enhanced in the second round (Fig. 4B), consistent with the TdT reaction and BrdUTP incorporation having a minimal effect on the antigenicity or distribution of albumin. These data suggest TUNEL can be performed after a MILAN immunostaining round entailing PC-antigen retrieval, immunofluorescence with MILAN buffers, 2-ME/SDS erasure, and that performing TUNEL did not influence immunostaining of a highly-expressed hepatocyte marker.

**Figure 4.**
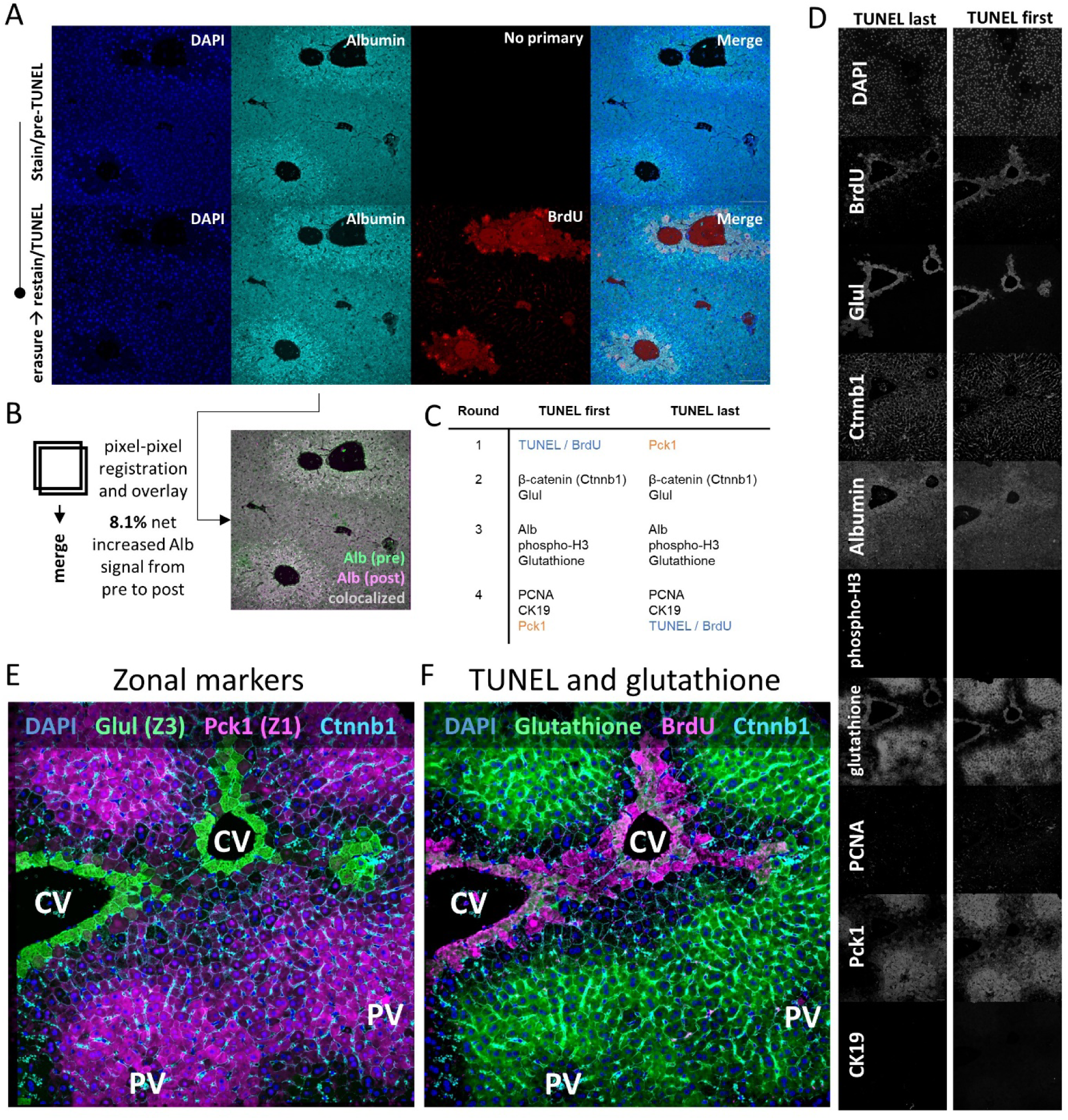
Versatile integration of TUNEL into a MILAN series enables spatial mapping of necrosis during acetaminophen toxicity. (A) MILAN immunofluorescence pre- (top row) and post TUNEL reaction (bottom row). (B) Mapping the effect of TUNEL on albumin staining by pixel-pixel image registration. The pre- and post-TUNEL albumin images (from Fig. 4A/cyan) were overlaid where white indicates equivalent brightness contributing from pre- and post, green indicates a bias toward higher intensity pre-TUNEL, and pink highlights higher intensity of albumin post-TUNEL. (C) Overview of MILAN staining rounds. (D) Single-channel immunofluorescence from each channel in a TUNEL-first slide versus a TUNEL-last slide in serial sections. (E) Spatially mapping the liver zones by MILAN immunofluorescence and image registration highlights central veins (Zone 3/Z3) proximal to Glul, and Zones 1-2 highlighted by Pck1 (TUNEL first series shown). Hepatocyte boundaries marked with Ctnnb1. (F) TUNEL staining as marked by BrdU in the same region, and MILAN immunofluorescence for glutathione. All scalebars are 100µm.

We then asked whether several rounds of MILAN immunofluorescence influenced TUNEL, and vice versa. In serial sections from 6h-APAP, TUNEL was performed either first or last (**Fig. 4C**), with otherwise four matched intervening rounds of immunofluorescence labeling proliferation (PCNA, pH3), synthetic function (albumin), hepatocyte membrane (β-catenin), cholangiocytes (CK19), and zonation markers (Zone 1-Pck1; Zone 3-Glul). Glutathione was also stained; glutathione depletion is thought to be a key step in acetaminophen-induced hepatocyte death. Encouragingly, TUNEL signal was preserved in the TUNEL-last slides, indicating that serial immunostaining and erasure rounds neither amplified nor reduced TUNEL reactivity (**Fig. 4D**). Immunostaining for the other 8 targets across three rounds appeared identical, indicating performing TUNEL first did not influence the antigenicity of subsequent rounds.

In addition to demonstrating versatile integration of TUNEL into a MILAN staining procedure, these data demonstrate high definition of the liver lobule architecture during acetaminophen hepatotoxicity (**Fig. 4E,F**). Glul staining demonstrates the edge of Zone 3, and correlates well with necrotic, TUNEL-positive cells. In contrast, glutathione is markedly depleted in Zone 3 consistent with prior observations that glutathione depletion is a key antecedent step in hepatocyte necrosis during acetaminophen-mediated injury. Taken together, these findings reveal that TUNEL can be deployed prior to MILAN, or up to four rounds into the iterative MILAN process, while preserving both TUNEL and immunostaining integrity. Harmonizing MILAN with TUNEL allowed single-cell resolution and spatial zonal contextualization of necrotic cell death during acetaminophen-induced hepatotoxicity.

### Cortical apoptosis and proliferation are increased in dexamethasone-treated adrenals

As noted above, costaining for the adrenal zona glomerulosa marker Dab2 and proliferative marker PCNA was negative in Click-iT treated serial sections. However, both were visualized with the PC-IH protocol, offering an opportunity to investigate proliferation and cell death spatially and in the same specimens.

The adrenal gland is structurally and functionally organized, featuring a capsule enveloping the adrenal cortex and an innermost adrenal medulla. In adult mice, the adrenal adult cortex is further organized into the outer, aldosterone-producing cells of the zona glomerulosa (zG) and the inner, glucocorticoid-producing cells of the zona fasciculata (zF). In subsequent MILAN rounds, we stained for additional structural markers, including Zo1, which was found to densely stain the capsule, and tyrosine hydroxylase, which highlighted the medulla (Fig. 5A-C). Coupled with Dab2, which marks the zG, the complete architectural structure of the zonated adrenal gland was specified spatially (**Fig. 5C**).

**Figure 5.**
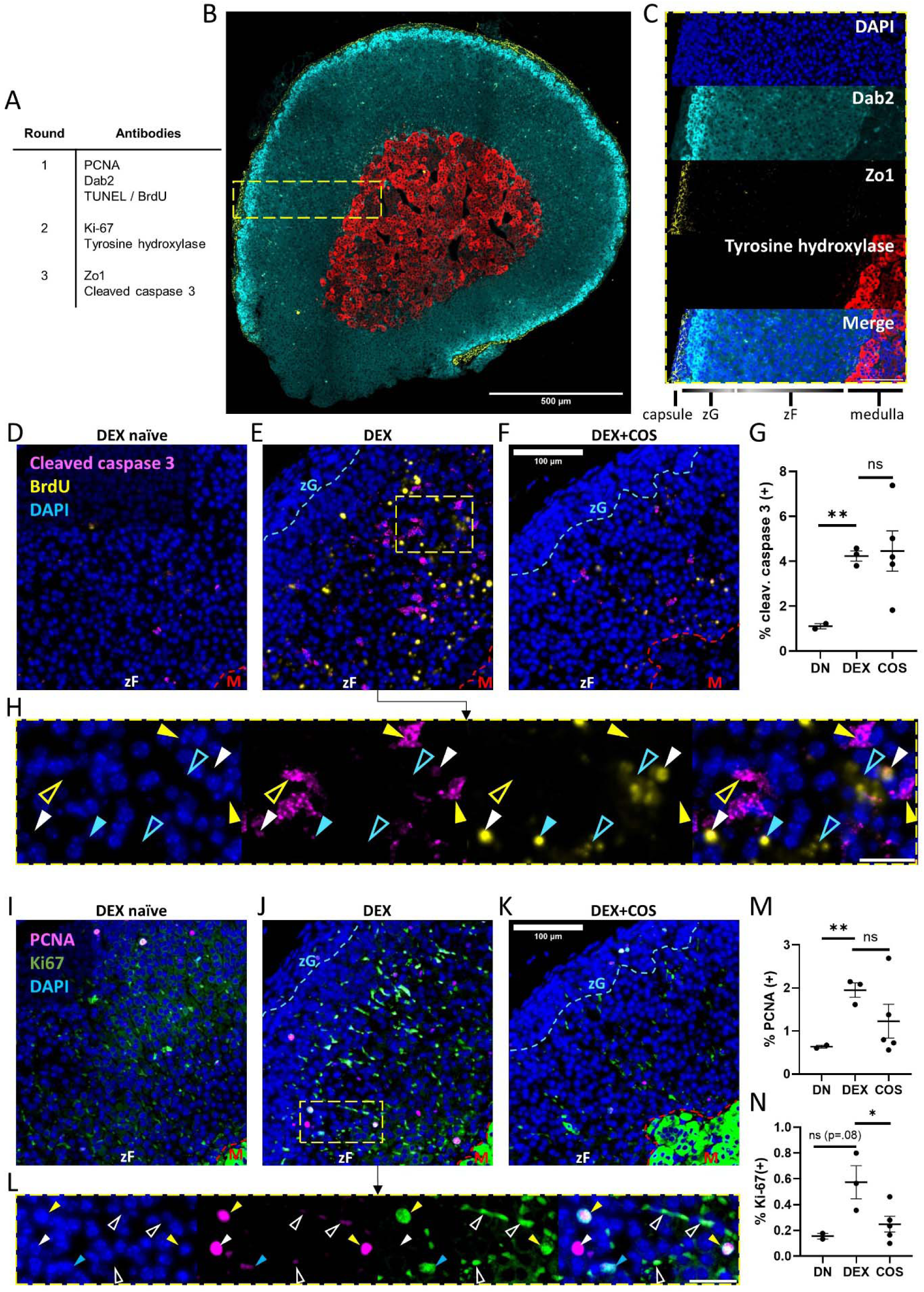
Dexamethasone increases adrenocortical apoptosis and proliferation. (A) MILAN staining series in adrenal glands with DEX, DEX+COS, or carrier/DMSO. (B) Spatially mapping the adrenal gland zones. (C) Inset of (B), scalebar is 100µm. (D-F) TUNEL and cleaved caspase 3 co-registration by harmonized TUNEL+MILAN with precise localization to the zF (zG-zF boundary (cyan) localized by Dab2, zF-medulla (M) boundary (red dashed) localized by Tyrosine hydroxylase), images representative. (G) Quantification of cleaved caspase 3 spots by treatment condition. (H) Inset of (E); scalebar is 25µm. Morphologically different patterns emerged: a fragmentation pattern consistent with apoptosis is noted by TUNEL/BrdU (open cyan arrows), a condensed TUNEL-only pattern (closed cyan arrow), colocalized TUNEL+cleaved caspase 3 (white arrow), DAPI colocalized cleaved caspase 3 (closed yellow arrow), and cleaved caspase 3 without DAPI colocalization (open yellow arrow). (I-K) Proliferation by MILAN immunofluorescence of PCNA and Ki-67 by adrenal condition. (L) Inset from (J) demonstrating variations of PCNA, Ki-67 and DAPI positivity to adjudicate true positive proliferating cells: PCNA+/Ki-67+/DAPI+ (yellow arrow), PCNA+/DAPI+ (closed white arrow), Ki-67+/DAPI+ (blue arrow), example Ki-67+/DAPI-background/autofluorescence not counted in proliferation quantification (open white arrow). Scalebar is 25µm. Quantification of PCNA (M) and Ki-67 (N) DAPI+ nuclei in the cortex.

Cleaved caspase 3 was added to corroborate detection of apoptosis, as TUNEL marks both apoptotic and necrotic cell death. TUNEL and cleaved caspase 3 were restricted to the zF (**Fig. 5D-F**). As with TUNEL, cleaved caspase 3 was markedly increased in DEX-treated glands (**Fig. 5G**). However, unlike TUNEL, there was no comparative rescue of cleaved caspase 3 in DEX+COS (**Fig. 5G**). TUNEL and cleaved caspase 3 did colocalize, but only in a minority of cells (**Fig. 5H**); in addition, the majority of both TUNEL and cleaved caspase 3 signal did not colocalize with DAPI signal (**Fig. 5H**), even in cases where the morphology was typical^21^ for a fragmented, apoptotic nucleus (**Fig. 5H**).

Examining proliferation (**Fig. 5I-L**), PCNA quantification of only the zF and zG layers disclosed upregulation of PCNA in DEX-treated samples compared to DN (**Fig. 5M**). Upregulation of PCNA was primarily in the zF in DEX-treated mice (**Fig. 5J**). ACTH rescue partially reduced proliferation, although this did not reach statistical significance. Because ACTH suppression, via dexamethasone, was strongly expected to inhibit rather than stimulate proliferation, we performed another round of MILAN using Ki-67. Many nuclei were double positive for PCNA and Ki-67 (**Fig. 5L**, yellow arrows), which elevated our confidence in the specificity of both antibodies. Ki-67 localization (**Fig. 5J**) and quantification corroborated cortical proliferation in DEX-treated samples and downregulation of Ki-67 in DEX+COS (**Fig. 5N**).

We conclude our harmonized PC-IH TUNEL-MILAN assay can be flexibly applied to multiple tissue types, enabling spatial contextualization of cell death. This harmonized protocol enables discovery of proliferation in the precise spatial adrenocortical region undergoing profound apoptosis.

## Discussion

TUNEL and immunofluorescence are two core techniques in the evaluation of tissue specimens with applications that range from characterizing model organisms to clinical diagnostics. Historically, both techniques consumed tissue, limiting the number and dimension of questions one could ask with individual specimens. While MILAN (and other cyclic staining methods) revolutionizes immunofluorescence by enabling many-fold erasure and restaining of FFPE specimens, TUNEL has heretofore required an individual specimen dedicated to its use. Here we have identified and optimized an erasure-compatible TUNEL protocol and harmonized this protocol with MILAN, discovering unexpected flexibility in the order of TUNEL in a MILAN staining series.

Prior to this work, consensus in the form of commercial assays and published protocols suggested that proK was required for optimal antigen retrieval in TUNEL. Indeed, TUNEL signal was greatest in proK-IH among the antigen retrieval strategies we tested in 6h-APAP livers. In addition, proK yielded a slightly more nuclear-enhanced pattern compared to protocols without proK. Despite this, pressure cooker antigen retrieval without proK was sufficient to retrieve characteristic staining of necrotic cells in the liver and apoptotic cells in the adrenal gland—the latter with equivalent signal-to-noise—indicating proK is not required for reliable detection of TUNEL signal. Others have reported that proK may be replaced with heat-mediated antigen retrieval or a pressure cooker^18^. Tsutsumi and Kamoshida found pressure cooker to be superior to proK in a variety of tissues including intestinal crypts, tingible body macrophages in necrotizing lymphadenitis and tonsillar germinal centers, invasive breast cancer, and in tuberculous granulomas^22^. They hypothesized that pressure cooker treatment may be particularly useful for elaborating old or over-fixated specimens; however, the specimens studied here were freshly processed and strictly fixed for 24-48 hours. Our study adds an additional advantage of consistency with pressure cooker treatment, in contrast to the high variability seen with Click-iT signal in both tissues.

ProK treatment was also broadly destructive of protein antigens in this study, with degradation or complete loss of all four proteins studied. This observation is of exceptional importance given thousands of published studies reporting co-immunofluorescence staining with conventional TUNEL protocols or commercial TUNEL kits, which generally employ proK pre-treatment. TUNEL protocol-induced loss of protein antigenicity has been reported previously^23,24^; however, much of the experimental evidence of altered antigenicity seems to be obscured in unpublished observations^23,25^. Commercial kits advise that proK time should be optimized in a tissue-specific manner for TUNEL reactivity, however, similar caution is clearly needed for protein antigenicity when conducting co-immunofluorescence. Moreover, proK treatment time may yield both false negative and false positivity TUNEL staining^25^, leading some to suggest there are no clear objective criteria to this optimization process^23^, now further complicated by the need to monitor for concomitant antigen loss when co-staining. Not only can antigen be lost by proK treatment, but our results demonstrate that localization of a protein can be aberrant, as seen with apparently nuclear-enhanced albumin expression. Such an artifact would be difficult to detect without performing a separate immunofluorescence experiment, which defeats much of the purpose of TUNEL co-staining in preserving a specimen. Collectively, the data presented here suggest extreme caution is warranted in the use of proK for conducting immunofluorescence.

The chief contribution of the present work is a harmonized protocol for TUNEL and MILAN-based multiplexed, iterative, tissue-preserving immunofluorescence. Pressure cooker treatment is compatible with, or possibly an improvement over proK for elaborating TUNEL signal and mitigates the primary incompatibility between traditional TUNEL and immunofluorescence protocols. We hypothesized that 2-ME/SDS might alter the antigenicity or distribution of TUNEL reactivity; however, at least across four cycles of immunostaining and erasure, TUNEL reactivity was well-preserved. The converse was also true: our revised TUNEL protocol could be inserted prior to MILAN immunofluorescence without affecting subsequent protein antigenicity.

We speculate a “TUNEL-first” PC-IH protocol may be compatible with other multiplex immunofluorescence approaches. In contrast, incorporating TUNEL late in a multiple staining and erasure protocol will require determining that other erasure procedures do not influence TUNEL (e.g., H_2_O_2_ in CyCIF^8^, guanidinum chloride / tris(2-carboxyethyl)phosphine^6^). Because H_2_O_2_ is known to induce single- and double-strand DNA breaks, CycIF may not be compatible with late-round TUNEL. In the MILAN-TUNEL protocol, further work is required to establish the limit, if any, of how many cycles into a MILAN protocol TUNEL may be incorporated without loss of TUNEL signal, with four cycles reported in this report.

This harmonized TUNEL-MILAN protocol enabled us to study two classic biological types of cell death, acetaminophen-induced hepatocellular necrosis, and dexamethasone-induced adrenocortical apoptosis. In APAP toxicity, cell death is thought to be restricted to Cyp2e1 expressing hepatocytes^26–28^, which reside preferentially in hepatocellular Zone 3 around central veins. Glutathione depletion is thought to be an essential step that mediates toxicity^29,30^, and indeed, multiplexed co-staining for zonal architectural coupled with glutathione and TUNEL signal shows strong anti-correlation between glutathione localization and necrotic cell death.

Our work also supports the well-established anti-apoptotic effects of ACTH, which is suppressed by glucocorticoids like dexamethasone, while raising new questions as to the mechanisms of adrenocortical proliferation in the absence of ACTH secretion. Wyllie and Kerr provided among the earliest morphologic descriptions of apoptosis in rat adrenals withdrawn from ACTH by either hypophysectomy or prednisolone injection, the latter of which was rescued with concurrent ACTH administration^15^. We similarly found that dexamethasone induced massive adrenocortical apoptosis, though concurrent cosyntropin treatment only partially attenuated TUNEL staining and had no effect on caspase-3 activation. There are several key differences between our and Wyllie and Kerr’s methodologies that may explain these results. First, our mice were treated for 4 days with dexamethasone and daily cosyntropin, a synthetic (1-24) ACTH analog thought to have the full potency of native ACTH^31^, versus 2 days with prednisolone and 3x/day purified ACTH. We are also actively investigating whether mice undergo tachyphylaxis to intermittent cosyntropin. This may explain why 4 days of combined dexamethasone and cosyntropin treatment reduced DNA fragmentation but not caspase-3 cleavage, the latter of which is a relatively early step in the apoptotic cascade. Tachyphylaxis to intermittent cosyntropin may also account for the more complete rescue of adrenocortical apoptosis defined by late-phase morphologic changes with two days of ACTH and prednisolone in Wyllie and Kerr’s studies compared to our work^15^.

The pro-proliferative adrenal effect in DEX-treated mice was unexpected given prior evidence that dexamethasone inhibits and ACTH stimulates adrenal proliferation. Short-term dexamethasone exposure (1 hour) reduced growth and proliferative pathways in both mouse adrenals and murine adrenocortical Y-1 cells^32^. Sustained exposure to both native and synthetic (1-24) ACTH stimulates adrenocortical hyperplasia *in vivo* after an initial period of hypertrophy^33^, though this has not been redemonstrated *in vitro*^33–35^. Hypophysectomy or corticosteroid treatment inhibits regeneration after bilateral adrenal enucleation but does not block compensatory adrenal growth after unilateral adrenalectomy, suggesting that ACTH is necessary for some but not all adrenal mitogenic programs^36–38^. Neural pathways, angiotensin II, IGF-1, FGF-2, endothelin 1, interleukin-6, VIP, NPY, and N-POMC have all been proposed as additional putative mediators of adrenocortical proliferation^39–43^. Our results suggest that zF cells are capable of ACTH-independent self-replication, perhaps in direct or indirect response to apoptosis of neighboring cells. Consistent with this, concurrent treatment with dexamethasone and cosyntropin attenuated both apoptosis and proliferation. Importantly, this unexpected finding would have been overlooked using a conventional TUNEL kit with PCNA co-staining, where PCNA signal was entirely abrogated, either by proK treatment or lack of pressure cooker retrieval. Further work is needed to understand the mechanisms and durability of this dexamethasone-induced proliferative response and its relationship to adrenocortical apoptosis.

In conclusion, TUNEL is an essential *in situ* tool for characterizing cell death, while the use of multiplexed immunofluorescence methods like MILAN are accelerating the potential for understanding spatially distributed phenomenon in biology and disease. We anticipate the unified method presented here will add a new and powerful dimension to highly multiplexed immunofluorescence methods and enable previously inaccessible insights into understanding mechanisms of cell death in a precise spatial context.

## Acknowledgments

The authors would like to thank Patrice Delaney for her constructive feedback on this manuscript.

## Funding

National Institutes of Health Grant T32DK007191 (MSS).

National Institutes of Health Grants R01DK090311, R01DK105198, R24OD017870 (WG).

National Institutes of Health Grant F32DK131795-02 (LSG).

Pediatric Endocrine Society Clinical Scholar Award (LSG).

This work was supported by a Boston Children’s Hospital Office of Faculty Development/Basic & Clinical Translational Research Executive Committees Faculty Career Development Fellowship (LSG).

## Methods

### Mice and treatment

We have previously investigated the role of acetaminophen on liver injury and regeneration^14^. Briefly, 3-month old male C57BL/6J mice were fasted for 12 hours, and then exposed to 300mg/kg acetaminophen (dissolved in 0.9% saline) at time zero by intraperitoneal injection versus saline carrier. Livers were harvested at 6, 24, 48, and 96 hours (n=3 per timepoint), and fixed in 10% neutral buffered formalin for two days, transferred to 70% ethanol, and then processed for paraffin embedding.

For studies of adrenocortical apoptosis, 8-12 week-old male C57BL/6J mice (n=2-5) were treated with either dexamethasone (DEX; 20 mcg/day=∼65 mg hydrocortisone equivalent/m^2^/day) or vehicle (DEX-naïve or DN) via drinking water. A third cohort was treated with dexamethasone as above and daily, intraperitoneal cosyntropin (ACTH1-24; 10 mcg/kg/day; DEX+COS) at 16:30 to align with the physiologic, circadian ACTH peak. Adrenals were harvested after 4 days of treatment, and adrenals were fixed overnight in 10% neutral buffered formalin, transferred to 70% ethanol, embedded in 4% low-melting agarose, and then processed for paraffin embedding.

### Slide preparation and deparaffinization

Staining was performed on 5-micron sections adhered to Superfrost Plus microscope slides (Fisher Scientific, cat # 22-037-246). Deparaffinization was conducted according to the manufacturer instructions for the commercial TUNEL assay with a series of baths starting with two 5-minute xylene baths followed by one 3-minute 50%:50% xylene-ethanol bath, one 5-minute ethanol bath, one 3-minute ethanol bath, one 3-minute 95% ethanol bath, one 3-minute 85% ethanol bath, one 3-minute 75% ethanol bath, one 3-minute 50% ethanol bath, one 5-minute 0.85% NaCl bath. Finally, slides were rehydrated in PBS for 10 minutes, and then sections were outlined with a hydrophobic barrier pen (cat # H-4000).

### Antigen retrieval methods

Four methods of antigen retrieval were investigated as pre-treatment for TUNEL and MILAN immunostaining, pressure cooker (PC) antigen retrieval, proK treatment, SDS/2-ME “stripping” buffer (STR), and no treatment (NT). Antigen retrieval followed immediately after PBS recovery from deparaffinization. For pressure cooking, slides were immersed in a staining jar containing Tris-EDTA buffer, pH=9 (Abcam ab93684) in a Biocare Medical Decloaking Chamber (Model No. DC2002) filled with 500mL of distilled H_2_O per manufacturer instructions. Slides were then cooled to 50°C (∼15-minute), washed in dH_2_0 for 5 minutes, and then transferred to PBS. For proK antigen retrieval, the commercial TUNEL kit proK (Thermo Fisher C10619) was used for both the Click-iT protocol and our adapted in-house proK protocol. Briefly, sections were covered with proK solution diluted 1:25 in pbs (as per kit protocol) and incubated for 15 minutes at room temperature. Sections were then washed twice in PBS for 5 minutes, fixed with fresh 4% paraformaldehyde, washed twice in PBS for 5 minutes, and rinsed in dH_2_0. For 2ME/SDS stripping buffer retrieval, deparaffinized slides were immersed in a slide mailer containing 10% sodium dodecyl sulfate (SDS) and 0.0104% 2-mercaptoethanol (2-ME)^9^, and the slide mailer was then incubated in a water bath at 56C for 90 minutes with intermittent shaking. Slides were then washed in TBS adjusted to pH 7.5 containing 0.02% tween 20, 0.1M sucrose, 0.001% sodium azide (MILAN wash buffer)^9^ for five washes (1 rinse, followed by 5-, 10-, 15-, and 30-minute washes). Finally, for no antigen retrieval, slides were washed in MILAN wash buffer (“permeabilization” given tween content of MILAN wash buffer) for 10 minutes immediately following deparaffinization and rehydration and then washed 3 times for 5 minutes in PBS.

### TUNEL procedure

Three TUNEL variants were considered, in combination with the antigen retrieval methods above: two commercial kits and an in-house assay. For commercial chromogenic TUNEL staining, the ApopTag Peroxidase In Situ Apoptosis Detection Kit (Millipore, S7100) kit was used according to manufacturer instructions. For fluorescence-based TUNEL detection, we used the Click-iT Plus TUNEL Assay with Alexa 647 kit (Thermo Fisher C10619) according to manufacturer instructions. The in-house (IH) TUNEL protocol was adapted from an Abcam, BrdU-based protocol (Abcam, ab66110) and classic references^2,4^. Briefly, following antigen retrieval, slides were placed back in PBS for 5 minutes. A-tailing was conducted using terminal transferase (New England Biolabs M0315S) according to manufacturer specifications in the presence of 10mM 5-Bromo-2’-Deoxyuridine 5’-Triphosphate (BrdUTP, Thermo Scientific, B21550), at 37°C in a dark humidified slide box for 90 minutes. Slides were then washed three times in PBS. The complete TUNEL procedure is notated by the antigen retrieval method followed by the TUNEL protocol (e.g., proK-Click-iT for proK antigen retrieval with the commercial click it assay, or PC-IH for pressure cooker antigen retrieval with in-house TUNEL). Five variants were considered: proK-Click-iT (abbreviated Click-iT), proK-IH, PC-IH, STR-IH, and NT-IH.

### DNAse treatment

For a positive control of double stranded DNA breaks, slides were treated with DNAse immediately following antigen retrieval. Slides were incubated for 30 minutes with 1 unit of DNase I (Roche, Cas. No. 9003-98-9) at room temperature, and then washed in dH_2_0.

### Immunofluorescence and erasure

For samples processed using the commercial kits, manufacturer instructions were followed explicitly for co-immunofluorescent labeling. Briefly, for co-immunofluorescence on proK-Click-iT slides, slides were washed in PBS, blocked for 1 hour in BSA, and then primary antibody incubation was conducted overnight in antibody diluent as per manufacturer recommendations. MILAN immunostaining^9^ was conducted as described in the MILAN version 5 protocol^10^. MILAN immunofluorescence comprised the same steps as conventional immunofluorescence, excepting the use of MILAN wash buffer in place of PBS for all wash steps, and MILAN antibody diluent instead of kit-described diluent. Erasure was also conducted as described^9,10^, using the same procedure as noted for STR antigen retrieval, except that stripping was conducted after a round of immunofluorescence. All primary and secondary antibody dilutions and incubation steps were otherwise the same between conventional immunofluorescence and MILAN immunofluorescence.

### Primary and secondary antibodies

Primary and secondary antibodies, and their dilutions for all experiments are listed in Table 1. Antibodies were diluted per protocol for co-staining with Click-iT TUNEL kits, or in MILAN antibody buffer^10^ otherwise.

**Table 1.**
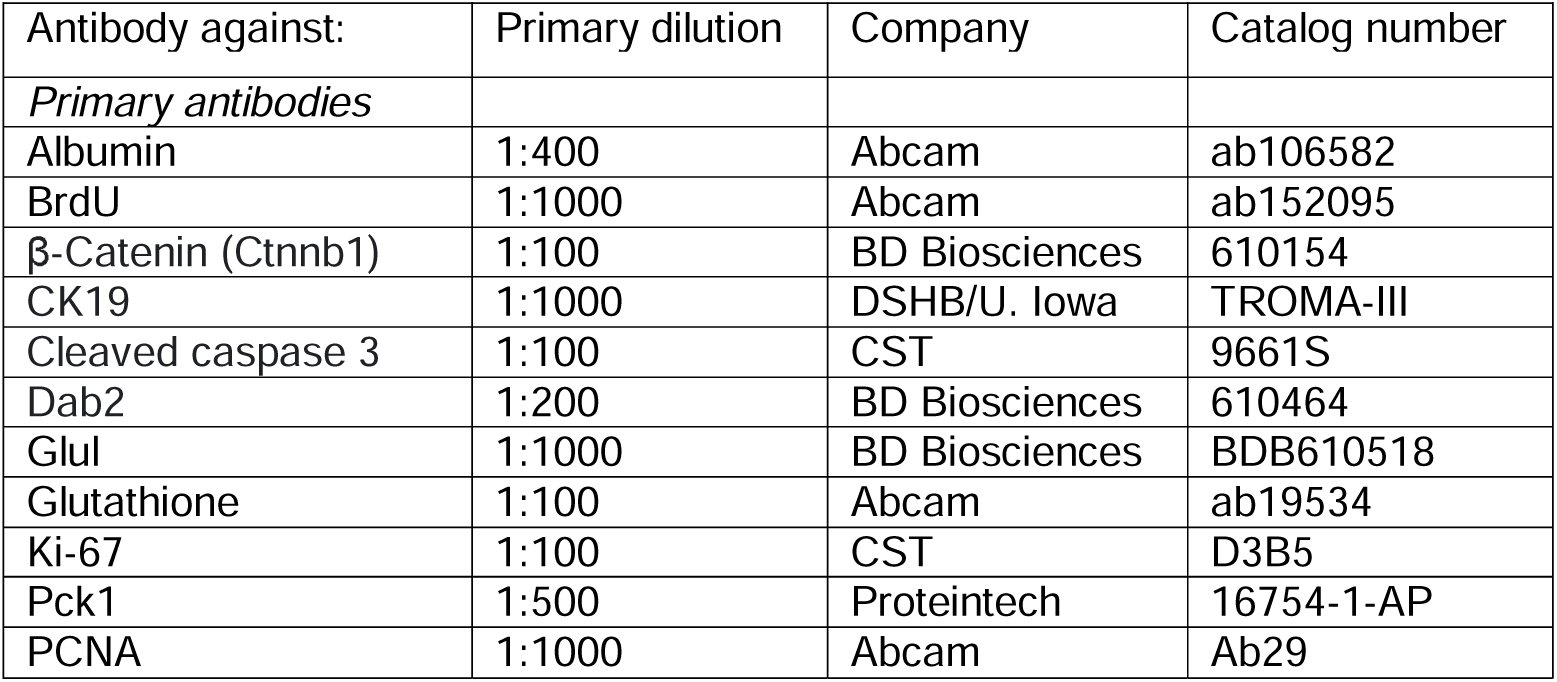
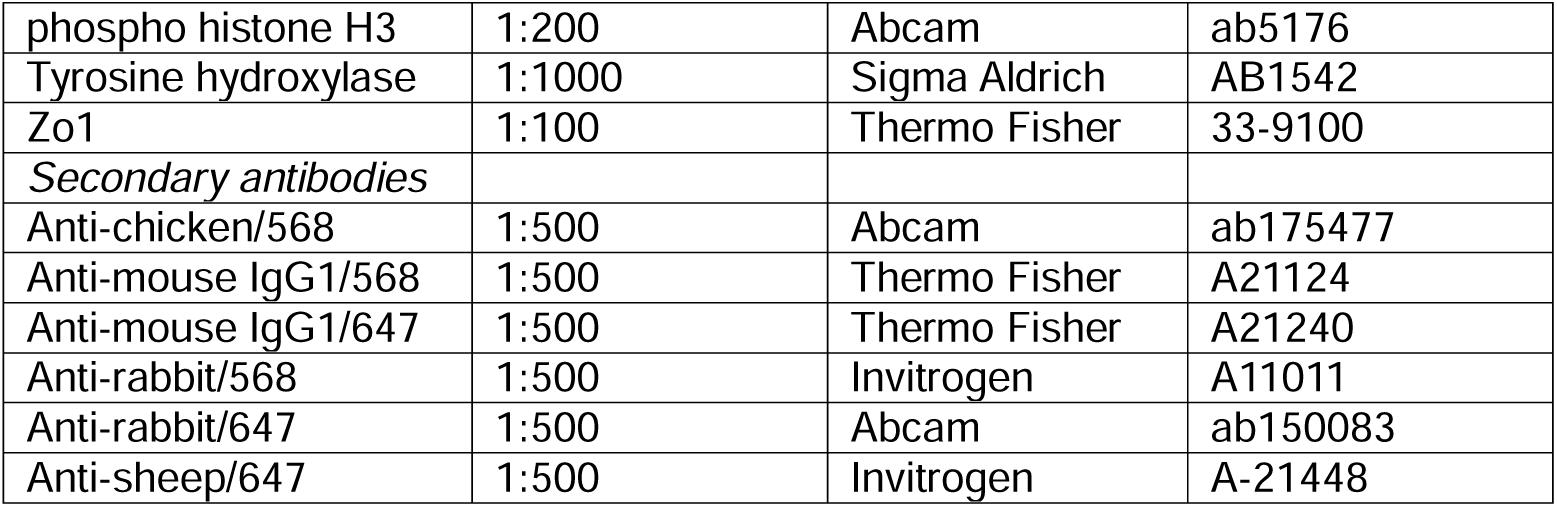
Antibodies and dilutions.

### Confocal microscopy

All confocal Microscopy was performed on a Nikon Ti2 microscope with a Yokogawa CSUW1 confocal unit, a Nikon Plan Apo Lambda 20x objective lens and a Zyla 4.2 PLUS sCMOS camera (ANDOR). All images were taken with identical laser strength, gain and exposure settings for the 405nM, 488nM, 568nM, and 647nM channels which were utilized by each image. Each image was acquired using the Large Image function in the NIS elements software. This function allowed us to take images in a 5×5 grid with 15% overlap allowing the images to be stitched together.

### Image analysis

For acetaminophen-exposed livers, the thickness of pericentral staining was measured in QuPATH. Images within each condition were blown out by moving the maximum contrast low and the minimum contrast up creating a thick band of positivity and little background staining, and then the brightness and contrast settings were applied to all images within the same condition. To calculate signal-to-noise, brightness of TUNEL-positive bands was measured across 4 images per biological replicate for each TUNEL method, and compared to the brightness of measured negative areas. Brightness was normalized by areas measured, thus, measurement was functionally the mean pixel signal to mean pixel noise. All measurements were obtained by one person, and statistically analyzed by another, to reduce analysis bias.

For TUNEL, nuclei, cleaved-caspase 3 and PCNA, spot counting was performed using Imaris (Oxford Instruments, Abingdon, United Kingdom). For each signal, a routine was generated that detects bright spots and tested on a few examples from each condition to ensure reasonable agreement by eye, tuning the threshold parameter to minimize both false positives and false negatives. Then, the routine was run using the same threshold for all images. All spots occurring within the medulla were excluded; this was done morphologically in the TUNEL quantification experiment comparing click-iT and PC-IH; subsequently, tyrosine hydroxylase staining of the medulla was used to exclude medullary cells. TUNEL positivity, PCNA positivity, cleaved-caspase 3 and Ki-67 positivity were calculated as the number of spots divided by cortical (non-medulla) DAPI spots, to account for differences in adrenal size or cuts. Signal to noise was calculated as mean TUNEL signal fluorescence divided by TUNEL negative DAPI signal fluorescence.

For β-catenin signaling intensity, the Find Maxima function in imageJ was used to select membrane edges (see. **Fig S3A**), although, when β-catenin staining failed this method likely measured autofluorescent spots rather than true staining. Thus, β-catenin quantification differences between PC and proK are likely conservative (see **Fig. S3B**). For albumin intensity measurements, 4 image fields per replicate were quantified, excluding portal/central veins, and normalizing for the areas measured.

## Conflict of interest statement

Authors declare they have no competing interests.

## Author contributions

Conceptualization: MSS

Methodology: MSS, LSG

Investigation: MSS, TM, LSG

Visualization: MSS, TM

Funding acquisition: WG

Project administration: MS, WG

Supervision: WG

Writing – original draft: MSS, TM, LSG, WG

Writing – review & editing: MSS, LSG, WG, JM

## Notes

### Competing Interest Statement

The authors have declared no competing interest.

