## Supplemental Information for "Harmonizing terminal deoxynucleotidyl transferase dUTP nick end labeling (TUNEL) with multiplexed iterative immunofluorescence enriches spatial contextualization of cell death"

**Supplemental Figures**

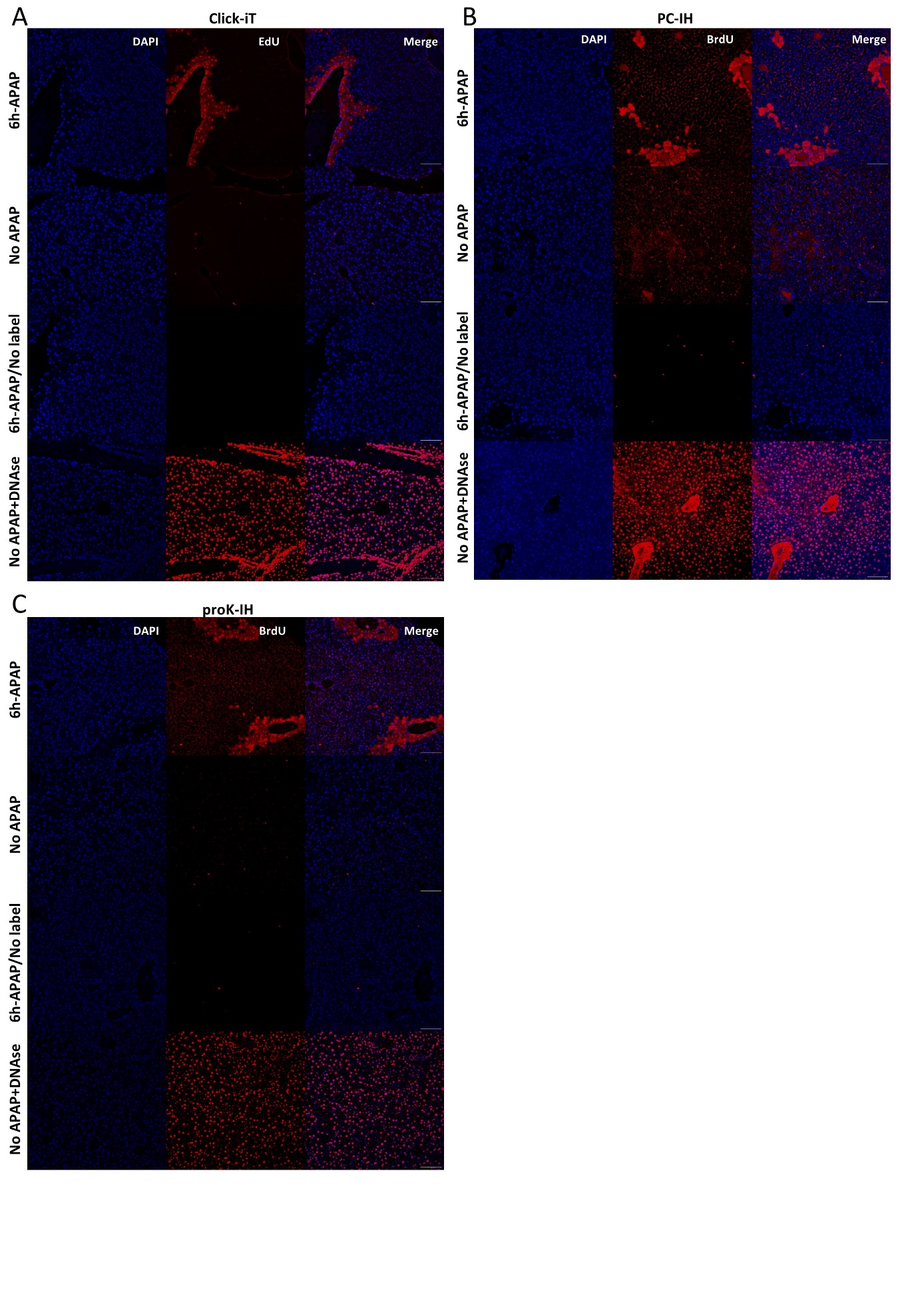

**Figure S1. Positive and negative controls for TUNEL protocol variants.** For Click-iT (A), PC-IH (B), and proK-IH (C), the top row corresponds to the standard TUNEL protocol performed on 6h-APAP murine liver samples. The second row corresponds to control mice that did not receive APAP, and serves as a biological negative control. The third row corresponds to a no-label (EdU for Click-iT; BrDU for -IH) negative control. The fourth row corresponds to a positive control whereby normal livers treated with DNase were processed with the full respective TUNEL protocol. All scalebars are 100µm.

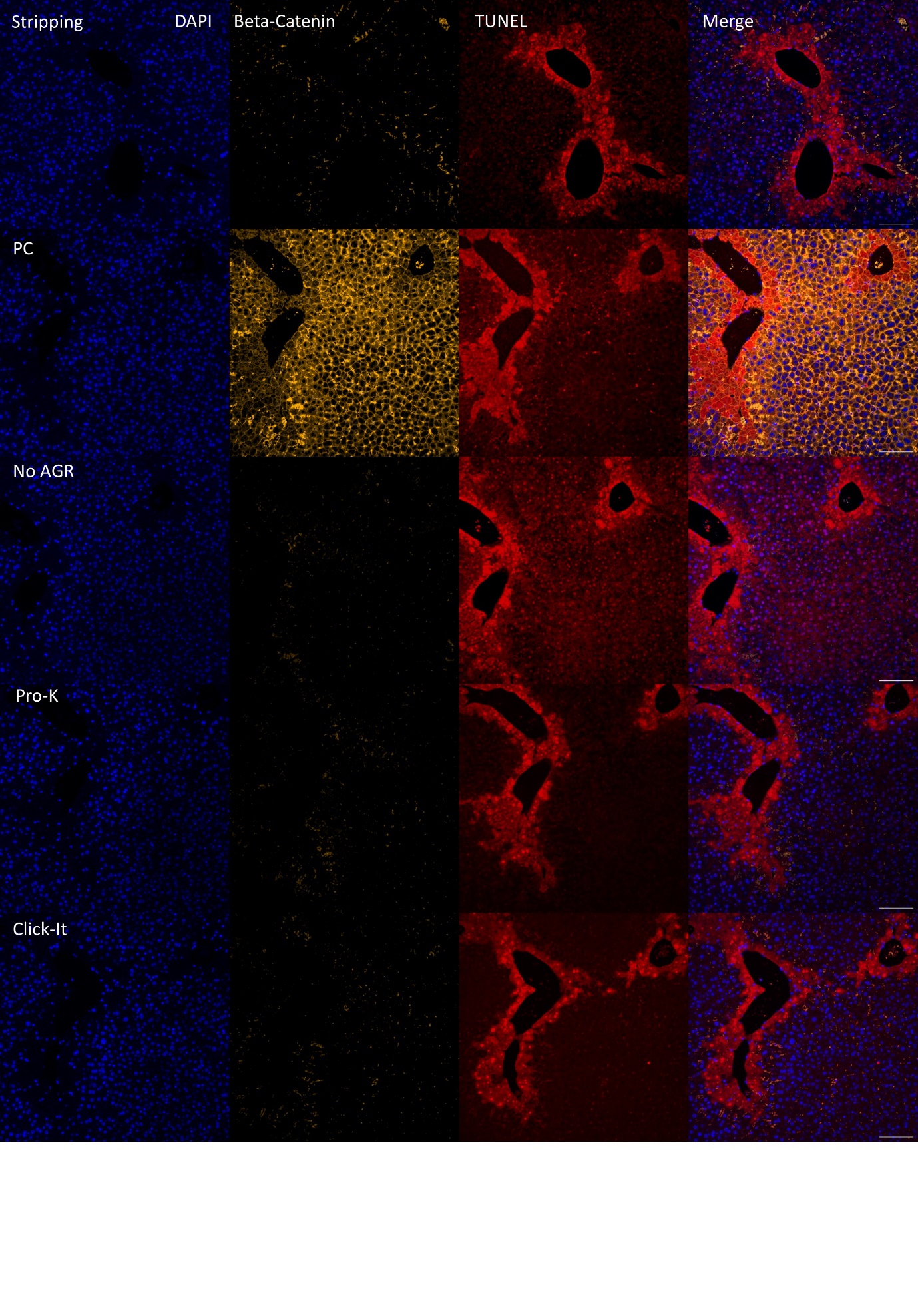

**Figure S2. TUNEL co-staining with β-catenin was detectable only with the pressure cooker antigen retrieval TUNEL version (PC-IH).** The antigen retrieval method is noted in the left-most column. All images are shown with the same quantitative brightness and contrast settings except for TUNEL signal in the Click-iT protocol, which was enhanced for morphologic comparison. All scalebars are 100µm.

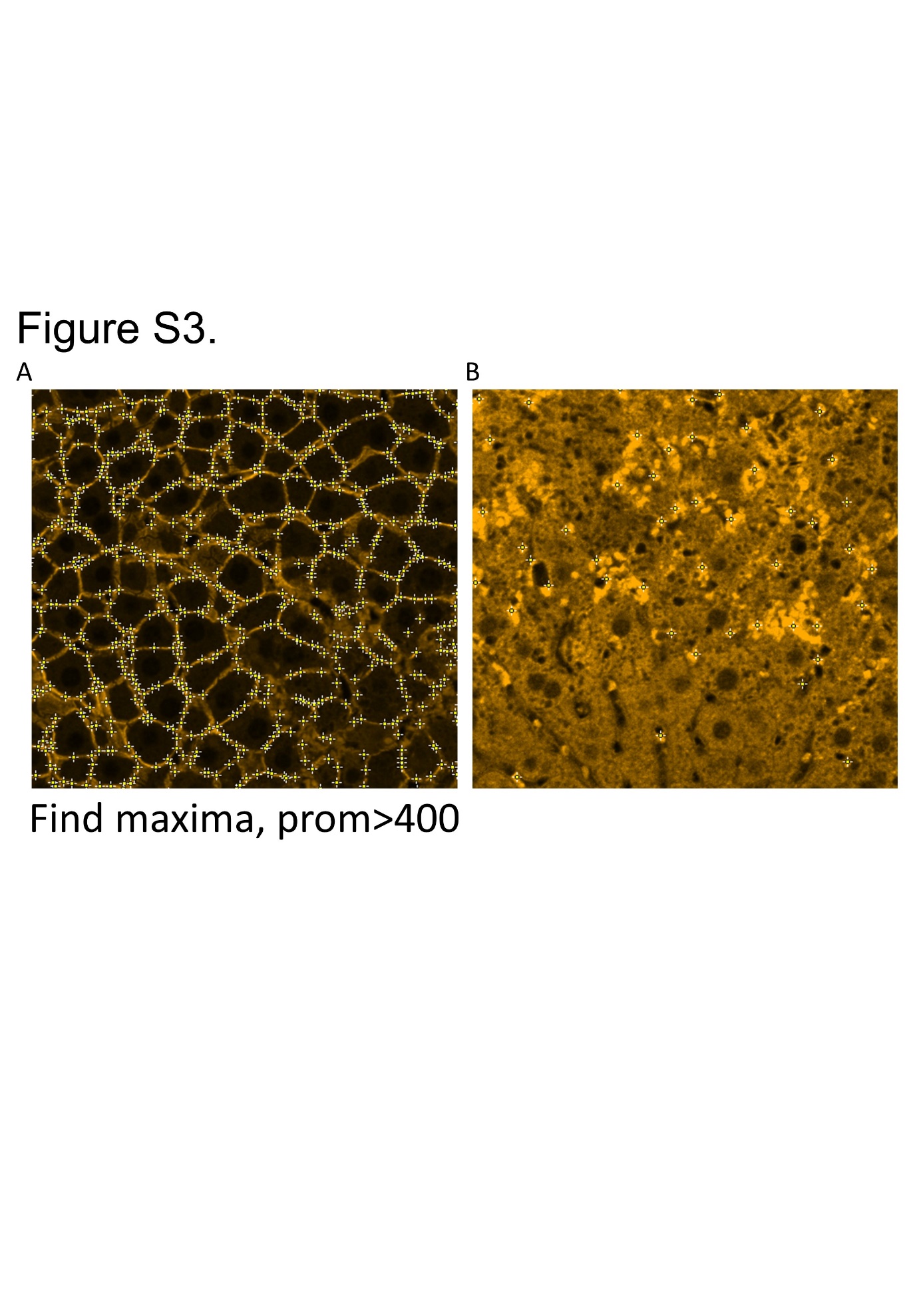

**Figure S3. Measuring β-catenin membrane intensity.** The Find Maxima function in imageJ was used to quantify mean β-catenin membrane intensity (A), setting the prominence to 400. In samples where β-catenin did not stain well, as in (B), the find maxima function serves as a conservative estimate.
